# Associations between EEG functional brain connectivity and a cognitive reserve proxy in healthy older adults

**DOI:** 10.1101/625608

**Authors:** Bahar Moezzi, Louise M. Lavrencic, Mitchell R. Goldsworthy, Scott Coussens, Hannah A.D. Keage

## Abstract

Cognitive reserve is a concept that explains individual differences in vulnerability to cognitive impairment due to age and dementia-related brain changes. Mechanisms underlying the cognitive reserve effect are poorly understood. We investigated associations between a comprehensive cognitive reserve proxy (Lifetime Experiences Questionnaire/LEQ) and functional connectivity of the prefrontal cortex across the whole scalp, covarying for the level of current cognitive functioning (Addenbrookes Cognitive Examination Revised/ACE-R), using multiblock parallel and orthogonalized partial least squares regression. EEG data were collected from 34 healthy older adults (63 to 83 years) in eyes-open and eyes-closed resting-states, and during 0-back and 1-back tasks. Functional connectivity was estimated using imaginary coherence in the theta and alpha frequency bands, as these bands have been heavily implicated in cognitive ageing, attention and executive function. We found three clusters of electrodes where the absolute values of the regression coefficient were above threshold when covarying for ACE-R: (1) a cluster approximating the right frontocentral region during the eyes-open condition in the theta band with seed electrodes approximating the left prefrontal cortex with positive associations of medium effect size; (2) a cluster approximating the right parietotemporal region during the 0-back task in the theta band with seed electrodes approximating the right prefrontal cortex with negative associations of medium to large effect sizes; and (3) a cluster approximating the occipitoparietal region in the eyes-closed condition in the alpha band with seed electrodes approximating the left prefrontal cortex with negative associations of medium effect size. These relationships between a cognitive reserve proxy and functional connectivity, within key networks and frequency bands associated with attention and executive function, may reflect greater neural capacity and efficiency.

## 1. Introduction

Cognitive reserve is a theoretical concept to explain individual differences in cognitive ageing and clinical dementia relative to neurodegenerative changes (Stern 2002; Stern 2012; Stern et al. 2008; Whalley et al. 2004; Wu et al. 2017). Those with higher cognitive reserve are less cognitively vulnerable to brain changes and have a lower risk of dementia in late life (Livingston et al. 2017; Stern 2002; Stern 2012; Whalley et al. 2004). Over two decades of research has consistently demonstrated a positive impact of cognitively stimulating activities, such as years of education and work complexity, on the risk of dementia and cognitive decline (Baumgart et al. 2015; Mortimer and Graves 1993; Schmand et al. 1997; Wang et al. 2002; Wilson et al. 2002; Xu et al. 2015; Zahodne et al. 2015). Cognitive reserve accrues over the lifespan via engagement in cognitively stimulating activities, which may be leisure or socially-based, along with education and occupation-related. Cognitive reserve cannot be measured directly, however, it is most commonly measured via proxies that capture these cognitively engaging activities over the lifespan.

A brain region that plays a critical role in the manifestation of cognitive reserve is the frontal cortex (Franzmeier et al. 2017; Solé-Padullés et al. 2009; Springer et al. 2005; Stern et al. 2008). In healthy young adults, higher scores on cognitive reserve proxies (indexed by two measures of IQ separately) were associated with increased activity in the right and left superior frontal gyri while performing a delayed item response task using fMRI (Stern et al. 2008). Right medial frontal gyrus activity during memory tasks in older adults has been shown to be positively correlated with years of education in older adults (Springer et al. 2005). However, during a visual encoding task, healthy older adults with higher values on a composite cognitive reserve measure exhibited decreased activity in the frontal lobe (Solé-Padullés et al. 2009). The difference in the direction of the effects in Springer et al. (2005) and Solé-Padullés et al. (2009) may be related to the methodological approaches (e.g. tasks differing in level of difficulty) and the analysis methods (i.e. univariate in Solé-Padullés et al. (2009) versus multivariate in Springer et al. (2005)). Notably, most of these fMRI studies did not control for current level of cognitive functioning, which is a critical step in identifying correlates of cognitive reserve as cognitive reserve proxies have been shown to positively associate with cognitive function in healthy older adults (Lavrencic, Churches, and Keage 2018; Opdebeeck, Martyr, and Clare 2016).

Another key issue within the cognitive reserve literature is how to reconcile conflicting reports of local frontal activity, i.e. some positive and some negative. Brains operate as networks (van Den Heuvel and Hulshoff Pol 2010) and therefore local activity only provides limited information about the brain mechanisms. Instead, functional connectivity between the frontal and distant cortical areas may better capture the neural mechanisms underlying cognitive reserve (Arenaza-Urquijo et al. 2013; Franzmeier et al. 2017; Stern et al. 2018). Using resting-state fMRI, higher global left frontal cortex connectivity has been associated with more years of education (a common proxy for cognitive reserve) in a prodromal Alzheimer’s disease sample (Franzmeier et al. 2017). In healthy older adults, years of education has been positively related to functional connectivity (using resting-state fMRI) between the anterior cingulate cortex and the hippocampus, inferior frontal lobe, posterior cingulate cortex and angular gyrus (Arenaza-Urquijo et al. 2013). In this study, Arenaza-Urquijo et al. (2013) assessed functional connectivity between brain areas that showed associations between years of education and both volume and metabolism, not across the whole brain. However, it is important to take into account the functional connectivity across the whole scalp because patterns of functional connectivity are not always related to gray matter volume or metabolism (Chételat et al. 2013; Ma et al. 2012). Further, hemispheric-dominant roles have been suggested for the frontal cortex in the expression of cognitive reserve (Franzmeier et al. 2017; Robertson 2014; Stern et al. 2008). Robertson (2014) proposed a right hemisphere frontoparietal network underlying cognitive reserve.

Conversely, left frontal (but not right) connectivity has been reported to be associated with higher scores on cognitive reserve proxies in normal and pathological ageing (Arenaza-Urquijo et al. 2013; Franzmeier et al. 2017).

Using fMRI data from participants aged 20-80 years performing 12 different cognitive tasks, a covariance functional connectivity pattern was identified whose expression correlated with a cognitive reserve proxy (IQ) (Stern et al. 2018). This task-invariant cognitive reserve network encompassed a range of brain regions including: (1) bilateral cerebellum, bilateral medial frontal gyrus and anterior cingulate, left superior temporal gyrus and right superior temporal gyrus (positive effects); and (2) right middle and inferior frontal gyri, right inferior parietal lobule, left precuneus and inferior parietal lobule, left middle and inferior frontal gyri (negative effects) (Stern et al. 2018). Notably, no resting-state was included. Patterns of functional connectivity showing associations with cognitive reserve proxies are known to differ during cognitive tasks compared with rest (Arenaza-Urquijo et al. 2013; Fleck et al. 2017; Franzmeier et al. 2017; Stern et al. 2018; Stern et al. 2008). Further, differences would be expected in different resting states, given functional brain networks in operation during the eyes-open resting-state are different from eyes-closed (Fleck et al. 2017; Xu et al. 2014), due to different attentional load and arousal levels (Hüfner et al. 2009).

Functional connectivity can be estimated in specific frequency bands using electroencephalography (EEG) recordings. Frequency bands of particular interest are theta and alpha, as they play a crucial role in cognitive processes (Başar and Güntekin 2012; Cavanagh and Frank 2014; Klimesch 1999; Klimesch 2012) and in cognitive ageing and dementia (Bhattacharya et al. 2013; Klimesch 1999). Alpha band power decreases, and theta power increases, from early to late adulthood (Dustman, Shearer, and Emmerson 1999); and compared to age-matched participants, people with dementia show higher theta and lower alpha power, and a shift in alpha activity from posterior to anterior recording sites (Başar and Güntekin 2012; Klimesch 1999). Using magnetoencephalography (MEG) data during a memory task, it was reported that in healthy older adults, cognitive reserve proxy scores (indexed by educational and occupational attainment) negatively correlated with theta and alpha functional connectivity (López et al. 2014). Conversely, eyes-closed resting-state EEG data from a sample of 90 cognitively normal adults (45 to 64 years) showed positive associations between higher cognitive reserve scores (years of education and verbal IQ) for theta and alpha connectivity (Fleck et al. 2017).

We aimed to identify whether theta and alpha functional connectivity is associated with a comprehensive cognitive reserve proxy, independent of current level of cognitive functioning. A thorough understanding of the mechanisms underlying the cognitive reserve effect requires investigation during tasks of various demand (Stern 2002; Stern 2009), so we tested a range of experimental conditions including eyes-open and eyes-closed resting-states and cognitive tasks of increasing difficulty (i.e. 0-back and 1-back tasks), investigating functional connectivity in theta and alpha bands for each of these conditions. Based on Fleck et al. (2017), we hypothesized that there would be a positive association between functional connectivity and the comprehensive measure of cognitive reserve in the theta and alpha bands at rest. Based on López et al. (2014) and Stern et al. (2018) we hypothesized that there would be a negative association between functional connectivity and the comprehensive cognitive reserve proxy during both 0 and 1-back tasks.

## 2. Materials and Methods

### 2.1. Participants

Thirty-four adults (63 to 83 years, mean age = 70.7 years, SD = 5.81 years, 14 males) were included in the study. Participants were followed-up from a larger study (n=115, see Lavrencic et al. (2016)) based on their scores on the Lifetime Experiences Questionnaire (LEQ), a comprehensive cognitive reserve proxy measure. The participants with LEQ at the extremes of the distribution were sampled first to ensure a spread of those with high and low cognitive reserve scores. Exclusion criteria for this study (part of a broader protocol) included: not a native English speaker, not strongly right-handed according to the Flinders Handedness Survey (Nicholls et al. 2013), history of alcohol or substance abuse or dependence in the past year, history of recreational drug use in the past month, history of any psychiatric disorder in the past 5 years, history of brain disorder or head injury, tendency to get headaches, diagnosis of a learning disability, clinical diagnosis of dementia, diagnosis of human immunodeficiency virus or Hepatitis C, hearing impairment or aid, uncorrected visual impairment, uncontrolled high blood pressure, history of cancer in the past 5 years (except skin or prostate cancer), taking medications targeting the central nervous system during the past month, taking medications that cause drowsiness, currently sleep deprived, or MRI contraindications.

### 2.2. Procedure

EEG data were recorded, and behavioural data collection (including cognitive reserve proxies, cognitive functioning and task performance scores) occurred during a three-hour testing session. All participants gave written informed consent in accordance with Australian National Health and Medical Research Council (NHMRC) Guidelines to participate in this study. Ethical approval for the study was granted by the University of South Australia’s Human Research Ethics Committee.

### 2.3. Measures

#### 2.3.1 Cognitive reserve: Lifetime of Experiences Questionnaire (LEQ)

The LEQ is a validated measure of cumulative mental activity measured across three life-stages: young adulthood, midlife, and late-life (Valenzuela and Sachdev 2007). Participants retrospectively recall information related to their educational, leisure, social and occupational history. The responses are weighted and combined into life-stage scores separately, and then are combined to generate a total LEQ score, with a higher score indicative of higher cognitive reserve.

#### 2.3.2. Cognitive functioning: Addenbrookes Cognitive Examination Revised (ACE-R)

ACE-R is a measure of cognitive functioning which can be utilized to screen for cognitive impairment and dementia (Larner 2007; Lonie, Tierney, and Ebmeier 2009; Mioshi et al. 2006). ACE-R includes five subscales measuring attention/orientation, memory, fluency, language, and visuospatial abilities. A Total ACE-R score (possible range 0-100) is summed from the subtests, with a higher score indicative of better cognitive functioning.

#### 2.3.3. EEG

We acquired two minutes of continuous resting-state EEG data with the eyes open and two minutes with the eyes closed. EEG data were also recorded during n-back tasks. The n-back tasks are suggested to test attentional control and executive function (Aalto et al. 2005; Kane et al. 2007; Owen et al. 2005). For the n-back tasks, stimuli consisted of five letters F, H, L, N, and T shown in white font on a black background, subtending a visual angle of 1.4° (width) by 1.5° (height). Each n-back task consisted of 40 target and 112 non-target stimuli. We presented the stimuli in a pseudo-randomized order one at a time for a duration of 500 ms. This was followed by a blank screen. The inter-stimulus interval jittered from 1200 ms to 1500 ms duration.

Participants underwent 0- and 1-back tasks. Notably a 2-back was run, however, over half of participants performed below chance, and we therefore excluded this task from analysis. For the 0-back, participants were asked to indicate when a target letter was presented (letters L or T, which were counterbalanced across participants). For the 1-back task, participants were asked to indicate to a target whenever the letter shown matched the one shown immediately before it. For all n-back tasks, target letters were responded to with one hand, and non-target letters with the other hand (counterbalanced across participants); the first three stimuli were not targets and there were no more than two of the same stimulus in a row. A practice trial was included before starting each n-back task. Response speed and accuracy were equally stressed.

For the duration of the EEG recording, participants were comfortably seated 60 cm in front of a 474×296 mm LED computer monitor. The EEG data were recorded using 64 Ag/AgCl electrodes embedded in a Quick-Cap (Compumedics), in accordance with the extended 10-20 international system (left: FP1, AF3, F7, F5, F3, F1, FT7, FC5, FC3, FC1, T7, C5, C3, C1, TP7, CP5, CP3, CP1, P7, P5, P3, P1, PO7, PO5, PO3, CB1, O1; midline: FPz, Fz, FCz, Cz, CPz, Pz, POz, Oz; and corresponding right channels). The ground was located between sites Fz and FPz, with the nose used as the reference. A Synamps II amplifier (Compumedics Neuroscan) was used to amplify the EEG using a band-pass filter of 0.05-200 Hz, and digitized at a sample rate of 1000 Hz using SCAN 4.5 acquisition software (Compumedics). The recorded data were stored on a computer for offline analysis.

#### 2.3.4. Task performance during n-back tasks: Inverse Efficiency Score (IES)

IES is a single behavioural outcome measure for task performance which accounts for potential criterion shifts and speed-accuracy trade-offs (Akhtar and Enns 1989). To compute IES, we extracted the reaction time and the total correct trials for all correct target and nontarget trials separately. IES was computed for each participant and condition (i.e., target or non-target) by dividing the mean reaction time by the proportion of correct responses, with a lower value indicative of a better performance. Mean IES values (for targets and non-targets) were used for analyses.

### 2.4. Data processing and analysis

EEG data were exported to MATLAB 9.0 (MathWorks, Inc., Natick, MA) for pre-processing and analysis. The signals were segmented into epochs of 1 s (for n-back: −0.2 s to 0.8 s with respect to each n-back letter stimulus). Channel baseline means were removed from the EEG dataset. Channels that were disconnected during recording or dominated by exogenous artefact noise were removed. Data were filtered using a hamming windowed sinc FIR (Finite Impulse Response) filter (1-45 Hz) and the EEG dataset was converted to average reference. We detected and excluded epochs contaminated by excessive noise by a procedure based on the identification of a threshold for the maximum allowed amplitude for the EEG signals >100 μV. Independent components analysis (ICA) artefact correction was used in order to correct for physiological artefacts (e.g. eye blinks and scalp muscle activity) (Delorme, Sejnowski, and Makeig 2007). Missing channels were interpolated using superfast spherical interpolation (Delorme and Makeig 2004).

We computed the power spectra for the EEG collected in the eyes-open and eye-closed resting-state conditions as well as the n-back tasks. To compute the power-spectra, we performed time-frequency analysis on EEG time series data over multiple trials using the multi-taper method based on Hanning tapers. The analysis windows were centred in each trial at 0.01 s intervals with three cycles per time window. We used the power spectra to set the range of the theta and alpha frequency bands. In line with Haegens et al. (2014) and Moretti et al. (2011), our alpha frequency was defined as the biggest local maximum within the extended range (5–14 Hz). The theta/alpha transition frequency (TF) was computed as the minimum power in the alpha frequency range so that TF < peak alpha frequency. Theta band was defined from 4 Hz to TF.

For EEG connectivity, we constructed the functional connectivity matrices for each participant using a highly conservative measure of connectivity, imaginary coherence, in theta and alpha frequency bands. Denote the *l*-th segment of the *i*-th time course by *x_i,l_* and its Fourier transform by *X_i,l_*. The cross spectral matrix is defined as

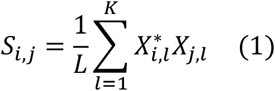

where (.)* denotes complex conjugation and *L* denotes the number of segments. Coherency between the *i*-th and *j*-th times series is defined as

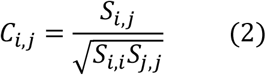

We computed the absolute value of imaginary coherence between each two electrodes *p* and *q* as

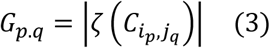

where *ζ*(.) denotes the imaginary part. Finally, matrix *G* is divided by its standard deviation using Jackknife method to produce a connectivity matrix.

We used parallel and orthogonalized partial least squares (PO-PLS) multiblock regression (Naes et al. 2013) to split the information in the input blocks connectivity and ACE-R into common and unique subspaces. PLS regression analysis is useful because it has the capacity to manage a higher number of independent variables than dependent variables without increased risk of Type I error and can handle non-orthogonal independent variables (Cramer and design 1993). Multiblock PO-PLS regression is useful because it has the capacity to handle different dimensionality of the blocks and is invariant to the relative weighting of the blocks (Krishnan et al. 2011; Naes et al. 2013).

We used PO-PLS multiblock regression to identify models unique to the functional connectivity covarying for ACE-R between (1) seed electrodes approximating left prefrontal cortex (LPFC: electrodes AF3, F3, F1, FP1) and all other electrodes and (2) seed electrodes approximating right prefrontal cortex (RPFC: electrodes AF4, F2, F4, FP2) and all other electrodes that maximally account for the variance in LEQ. We considered LEQ as a continuous variable in the analysis. Separate multiblock regression PO-PLS models were developed for the theta and alpha frequency bands.

A threshold of 0.3 reflecting a minimum value for a medium effect size was used for regression coefficients (standardized betas) to include in the PO-PLS model. In accordance with previous PLS studies, we utilized the first component for all PO-PLS models (Hordacre et al. 2017; Hordacre, Moezzi, and Ridding 2018). We computed percentage explained variance in LEQ uniquely by the functional connectivity in order to find out which conditions and frequency bands provide important contributions. In each PO-PLS model, we identified clusters of electrodes with at least three adjacent electrodes in space with regression coefficients above threshold. For power spectra and connectivity computations we used FieldTrip which is a MATLAB software toolbox for EEG and MEG analysis (Oostenveld et al. 2011). For PO-PLS analysis we used the Nofima multiblock regression by PO-PLS MATLAB toolbox (Naes et al. 2013).

Statistical analysis was performed using MATLAB 9.3 (MathWorks, Inc., Natick, MA). A median split on subjects’ LEQ scores was performed to classify them as either low cognitive reserve or high cognitive reserve for the purposes of assessing power across the EEG frequency spectra (otherwise, LEQ was always included as a continuous variable in models). Power at each 1 Hz frequency was compared between low cognitive reserve and high cognitive reserve participants by using nonparametric Mann-Whitney *U*-test (FDR correction for multiple comparison, *q* < .05). These analyses were done to ensure that frequency band windows did not shift relative to LEQ. Association between ACE-R with LEQ and between IES with LEQ was separately investigated with Spearman’s rank correlations (FDR correction, *q* < .05).

## 3. Results

### 3.1. EEG power

To set the appropriate range of frequencies for theta and alpha bands and to investigate whether frequency band windows varied with LEQ, we plotted the power spectra for all resting and task conditions – see Figure 1. This Figure shows that frequency bands 4-6 Hz for theta and 7-14 Hz for alpha (standard cut-offs) are suitable for our further analysis and the peak frequencies do not appear to differ between participants with high and low LEQ scores, i.e. we qualitatively observed no shift. We also observed that participants with higher LEQ scores using a median split showed significantly higher theta and alpha power.

**Figure 1.**
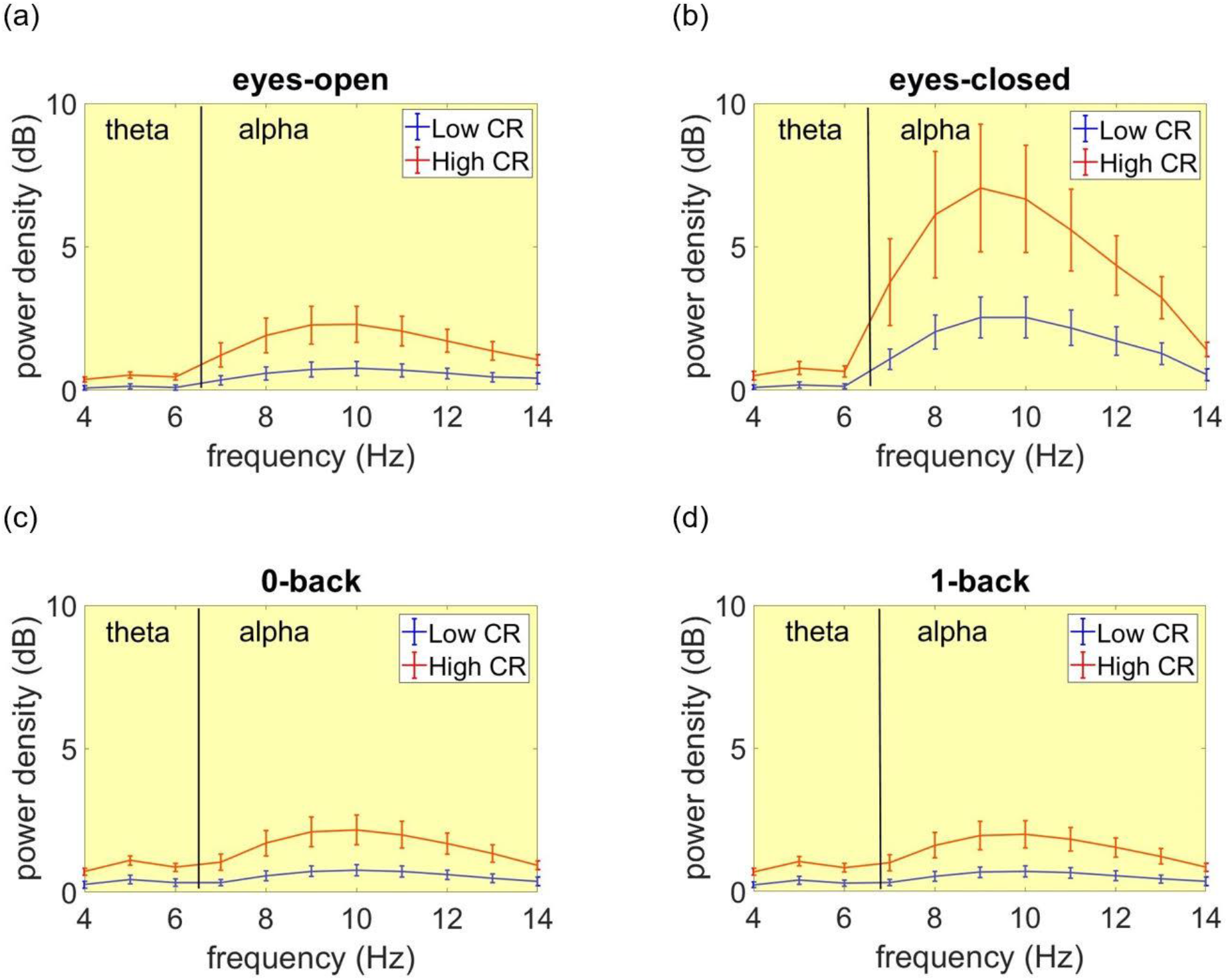
Comparison of power for subjects with low and high LEQ scores (median split) using a Mann-Whitney U-tests (FDR correction for multiple comparison, *q* < .05). Yellow shade: significant. (a) resting-state eyes-open, (b) resting-state eyes-closed, (c) 0-back task, and (d) 1-back task conditions. Error bars are SE. The black vertical lines denote the boundary of theta and alpha bands.

### 3.2. Associations between EEG functional connectivity and LEQ

#### 3.2.1. Resting-state

The multiblock PO-PLS models with seed electrodes approximating either RPFC or LPFC identified clusters of electrodes where the absolute value of regression coefficient was above threshold. In the eyes-closed resting-state condition, a cluster approximating the occipitoparietal region in the alpha band was found, with seed electrodes approximating LPFC when covarying for ACE-R scores (Figure 2), indicating lower alpha functional connectivity between electrodes approximating LPFC and occipitoparietal regions for those with higher LEQ scores. This pattern of effects was consistent when excluding the ACE-R as a covariate; however, the effects did not reach our threshold (see Supplementary Materials). No effects met threshold in the theta band (Supplementary Materials).

**Figure 2.**
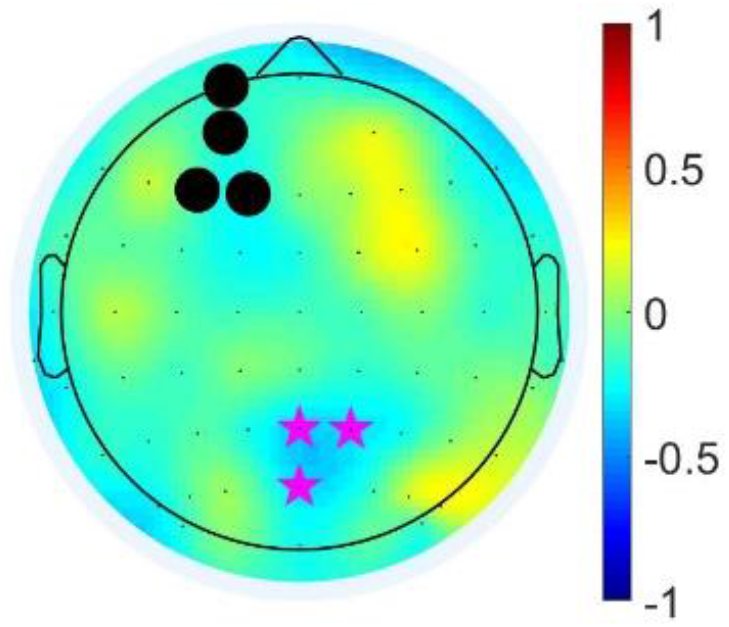
Topographic plots of regression coefficients from the parallel and orthogonalized partial least squares multiblock regression model with seed electrodes approximating LPFC in eyes-closed resting-state condition in alpha frequency band covarying for ACE-R. The seed electrodes are shown with filled black circles, and electrodes identified as being in a cluster are marked with magenta stars.

In the eyes-open resting-state condition, we found a cluster approximating the right frontocentral region in the theta band with seed electrodes approximating LPFC showing positive association between connectivity and LEQ scores when including ACE-R scores as a covariate (see Figure 3), and when the ACE-R was not included (although the effects did not result in any clusters of electrodes; see Supplementary Materials). No effects met threshold in the alpha band (Supplementary Materials).

**Figure 3.**
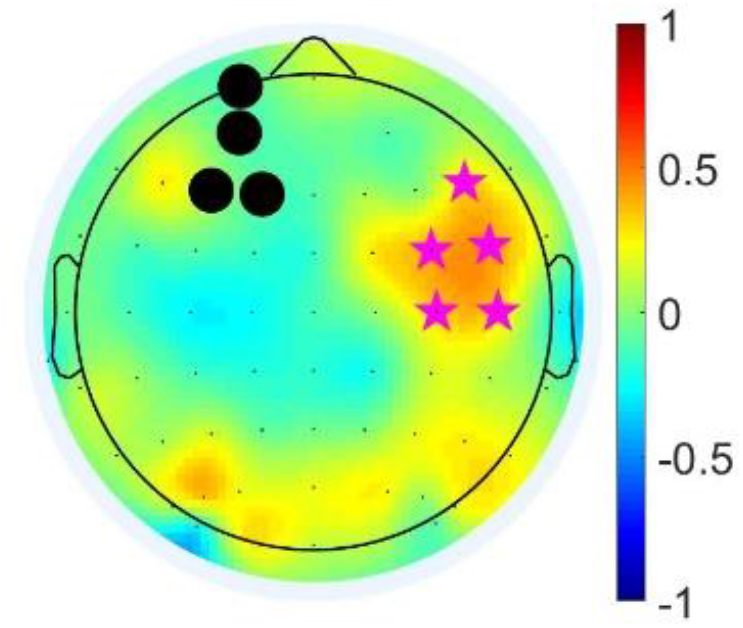
Topographic plots of regression coefficients from the parallel and orthogonalized partial least squares multiblock regression model with seed electrodes approximating LPFC in eyes-open resting-state condition in theta frequency band covarying for ACE-R. The seed electrodes are shown with filled black circles, and electrodes identified as being in a cluster are marked with magenta stars.

#### 3.2.2. N-back

Higher LEQ scores were associated with lower theta functional connectivity during 0-back between electrodes approximating RPFC and right parieto-temporal regions, when covarying for ACE-R scores (Figure 4) and when ACE-R was not included in the model (Supplementary Materials). No clusters were found for the alpha band (Supplementary Materials).

**Figure 4.**
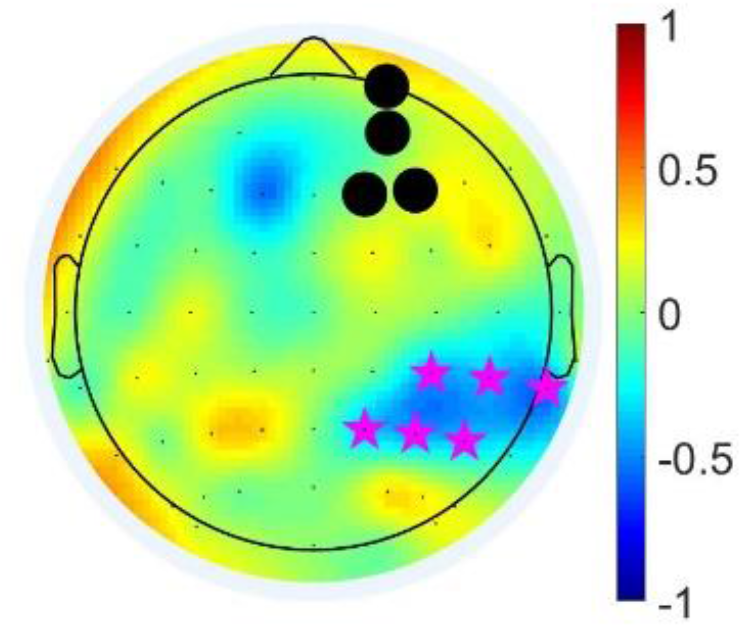
Topographic plots of regression coefficients from the parallel and orthogonalized partial least squares multiblock regression model with seed electrodes approximating RPFC in 0-back condition in theta frequency band when covarying with ACE-R. The seed electrodes are shown with filled black circles, and electrodes identified as being in a cluster are marked with magenta stars.

No clusters for either alpha or theta band were found for the 1-back condition (see Supplementary materials). Regression coefficients of clusters meeting threshold for each condition had medium to large effect sizes (by definition, all above r = 0.3).

### 3.3 Associations between cognitive functioning and LEQ

We further investigated the association of IES with LEQ. We observed that LEQ scores were not associated with IES in any of the task conditions (0-back: *rho* = −.20, *p* = .23; 1-back: *rho* = .27, *p* = .23).

We further investigated the association of ACE-R with LEQ. We observed that ACE-R had a significant positive correlation with LEQ scores (*rho* = .56, *p* = .001).

## 4. Discussion

This study aimed to investigate whether a comprehensive proxy measure of cognitive reserve was associated with theta and alpha functional connectivity at rest and during a cognitive task. We found different associations with cognitive reserve scores across these conditions. Higher cognitive reserve proxy scores were associated with lower functional connectivity for eyes-closed resting-state and the 0-back task conditions, and higher functional connectivity for eyes-open resting state, independent of cognitive function. There were no associations with functional connectivity for the 1-back task. Higher cognitive reserve scores were also associated with greater power in the alpha and theta bands.

In the eyes-closed resting-state condition we observed negative correlations between functional connectivity and cognitive reserve scores in the alpha band. In contrast, the associations between functional connectivity and cognitive reserve scores were positive in the theta band for the eyes-open condition. The shift from alpha to theta band during eyes-closed to eyes-open resting state suggests that the eyes-closed condition reflects a ‘true’ resting-state compared to eyes-open. The lower functional connectivity in alpha band for those with higher cognitive reserve scores during eyes-closed suggests that these individuals engage attentional networks to lesser degree at rest, reflecting more efficient neural networks.

Efficiency has been characterised by the rate of change in task activation with increasing task demand, so that a more efficient network will show less activation to produce the same (or better) level of performance, particularly when task demand is low (Stern 2002; Stern 2009; Stern 2012). This theoryhas been supported by previous studies demonstrating an inverse relationship between cognitive reserve (indicated by proxy measures) and functional connectivity during rest in various functional brain networks including the default mode and dorsal attention networks (Bastin et al. 2012). That there was greater functional connectivity in theta band during eyes-open for those with higher cognitive reserve scores does not fit with the efficiency hypothesis, but instead suggests that those with higher cognitive reserve scores may have been more actively engaged and attending during the task. As the eyes-open resting-state condition required participants to attend to a fixation cross, it is possible that the greater activation reflects self-imposed task demands (e.g., fully attending during fixation) in those with higher cognitive reserve scores. This is supported by the fact that it was functional connectivity in theta, and not alpha band, which was different for those with higher and lower cognitive reserve scores during the eyes-open rest. Theta band has been implicated in cognitive scaffolding, supporting the idea that those with higher cognitive reserve scores were cognitively engaged in the eyes-open resting state condition (Cooper et al. 2019; Reuter-Lorenz and Park 2014).

The negative correlations between functional connectivity and cognitive reserve scores in the eyes-closed condition, are in the opposite direction to the positive associations reported in previous literature between eyes-closed resting-state functional connectivity and cognitive reserve proxies in older adults (Arenaza-Urquijo et al. 2013; Fleck et al. 2017); although there are limitations with comparing fMRI and EEG findings. This directional difference could be driven by (1) the age of our sample (Vysata et al. 2014) as age range was 63 to 83 years in our sample and 45 to 64 years in Fleck et al. (2017) and (2), that our connectivity estimates were specific to the alpha frequency, facilitated by EEG recordings instead of fMRI in Arenaza-Urquijo et al. (2013). The pattern of positive associations seen in the eyes-open condition, however, are in general agreement with positive associations previously reported between eyes-open resting-state functional connectivity and cognitive reserve proxies in older adults (Franzmeier et al. 2017).

We observed negative correlations between functional connectivity of the electrodes approximating the right frontoparietotemporal regions and cognitive reserve proxy scores in the theta band for 0-back task. In humans, distributed theta oscillations have been reported to have particular relevance to attention (Sauseng et al. 2007). Theta frontoparietal connectivity is suggested to reflect the continuous cognitive processing (attention and executive control) needed to perform a task (Cooper et al. 2015; Corbetta and Shulman 2002; Coull, Frackowiak, and Frith 1998; Mizuhara and Yamaguchi 2007; Paus et al. 1997; Pesonen, Hämäläinen, and Krause 2007). Using positron-emission tomography (PET) and EEG recordings in healthy participants during an auditory vigilance task, it was reported that a right hemispheric theta frontoparietal network is engaged during attentionally demanding tasks (Coull, Frackowiak, and Frith 1998; Paus et al. 1997). Functional cortical circuits for executive functions have also been shown to be indexed by theta frontoparietal functional connectivity using simultaneous fMRI and EEG (Mizuhara and Yamaguchi 2007). Our findings are in line with reported negative associations between frontoparietal functional connectivity and cognitive reserve proxies during task by Stern et al. (2018), and a MEG study reporting negative associations between average theta functional connectivity across the scalp and cognitive reserve proxies (López et al. 2014). Unlike López et al. (2014), we assessed a task with increasing levels of difficulty, and have shown that the effects are only at low cognitive load.

Theta frontoparietal pattern is reflective of neural efficiency for those with higher cognitive reserve scores (for a review see Cabeza et al. (2018)). Our results may therefore reflect more efficient attention and executive function networks during low, but not high, task demands in those who score higher on a cognitive reserve proxy; as right theta frontoparietal functional connectivity is suggested to reflect attention and executive control (Cooper et al. 2015; Corbetta and Shulman 2002; Coull, Frackowiak, and Frith 1998; Klimesch 1999; Klimesch et al. 2005; Mizuhara and Yamaguchi 2007; Paus et al. 1997; Pesonen, Hämäläinen, and Krause 2007; Sauseng et al. 2007). We did not see the same results for the higher task demand condition (1-back). It is possible that no effects were seen for the 1-back task as the task was difficult for all participants regardless of cognitive reserve score, resulting in no functional connectivity differences. The right predominance seen in the 0-back task fits with the theory by Robertson (2014), which draws on empirical data from studies of arousal, sustained attention, response to novelty and awareness (Bellgrove et al. 2006; Greene et al. 2009; Grefkes et al. 2009; Naghavi and Nyberg 2005).

In line with previous studies we showed that individuals with higher cognitive reserve proxy scores exhibit higher theta and alpha power (Jaušovec 2000; Thatcher, North, and Biver 2005); and extended these findings by using a more comprehensive cognitive reserve proxy measure. Individuals with higher IQ have been reported to show higher alpha power at rest and during tasks (Jaušovec 2000). Power has been reported to be positively correlated with IQ in the alpha, theta and beta frequency bands in a resting eyes-closed EEG study from 442 individuals aged 5–52 years (Thatcher, North, and Biver 2005). Lower alpha power reflects a lower dominant oscillatory network activity modulated by thalamocortical and corticocortical interactions and hence shows a reduced global attentive readiness (Brunia 1999; Cantero et al. 2009; Da Silva et al. 1973; Klimesch 1999; Nunez, Wingeier, and Silberstein 2001). Increased electrical currents and consequently power density are suggested to be positively related with cognitive reserve proxies (Pellerin et al. 2007; Kapogiannis and Mattson 2011; Stranahan and Mattson 2012). It has been suggested that the brain cell energy metabolism, and hence the brain electrical activity (Pellerin et al. 2007), and cognitive reserve are linked (Kapogiannis and Mattson 2011; Stranahan and Mattson 2012).

A potential application of our study is in identifying the target brain regions and frequency bands in EEG neurofeedback training. EEG neurofeedback includes a brain-computer interface that enables users to learn to voluntarily control their cortical oscillations by obtaining online feedback from their EEG (Spilker et al. 1969; Taya et al. 2015). It is suggested that neurofeedback training can be used to modify functional brain network interactions (Haller et al. 2013; Horovitz, Berman, and Hallett 2010; Paret et al. 2016). If EEG-based functional connectivity neurofeedback training is to be used to enhance/maintain cognitive performance in healthy older adults, it may be important to identify the target connections that are most strongly associated with cognitive reserve proxy measures. Our results may have implications in designing effective neurofeedback rehabilitation strategies to enhance cognitive ageing.

We acknowledge that EEG suffers from poor spatial resolution and signals recorded at the scalp can be affected by volume conduction (Bastos and Schoffelen 2016). We have mitigated the volume conduction effects by using a highly conservative measure of functional connectivity (i.e. imaginary coherence). However, this approach may not thoroughly remove effects of volume conduction. Caution is required when suggesting generators of neural signal recorded with EEG surface electrodes. When explaining our results, we cautiously referred to electrodes clusters as approximating cortical regions as it is not clear where recorded surface signals are generated. Other limitations include our small sample size for a neuroimaging study and that we only considered one task type and did not consider other cognitive domains, albeit the task did include conditions with differing levels of cognitive load.

We observed effects for our eyes-closed resting condition (left lateralized; alpha band), eyes-open resting condition (right lateralized; theta band) and 0-back working memory task (right lateralized; theta band). Cognitive reserve appears to manifest as greater efficiency of frontoparietal and frontocentral networks during low demand states. Our findings may suggest greater efficiency to use key networks associated with attention and executive function in individuals with higher cognitive reserve.

## Acknowledgements

HADK is supported by a NHMRC Boosting Dementia Research Leadership Fellowship (GNT1135676). MRG is supported by a NHMRC-ARC Dementia Research Development Fellowship (1102272). LL is supported by a Serpentine Foundation Fellowship. We would like to thank Ingrid Måge of Nofima for helpful discussions. BM would like to thank Luke Hallam for helpful discussions.

## Supplementary materials

**Table 1.**
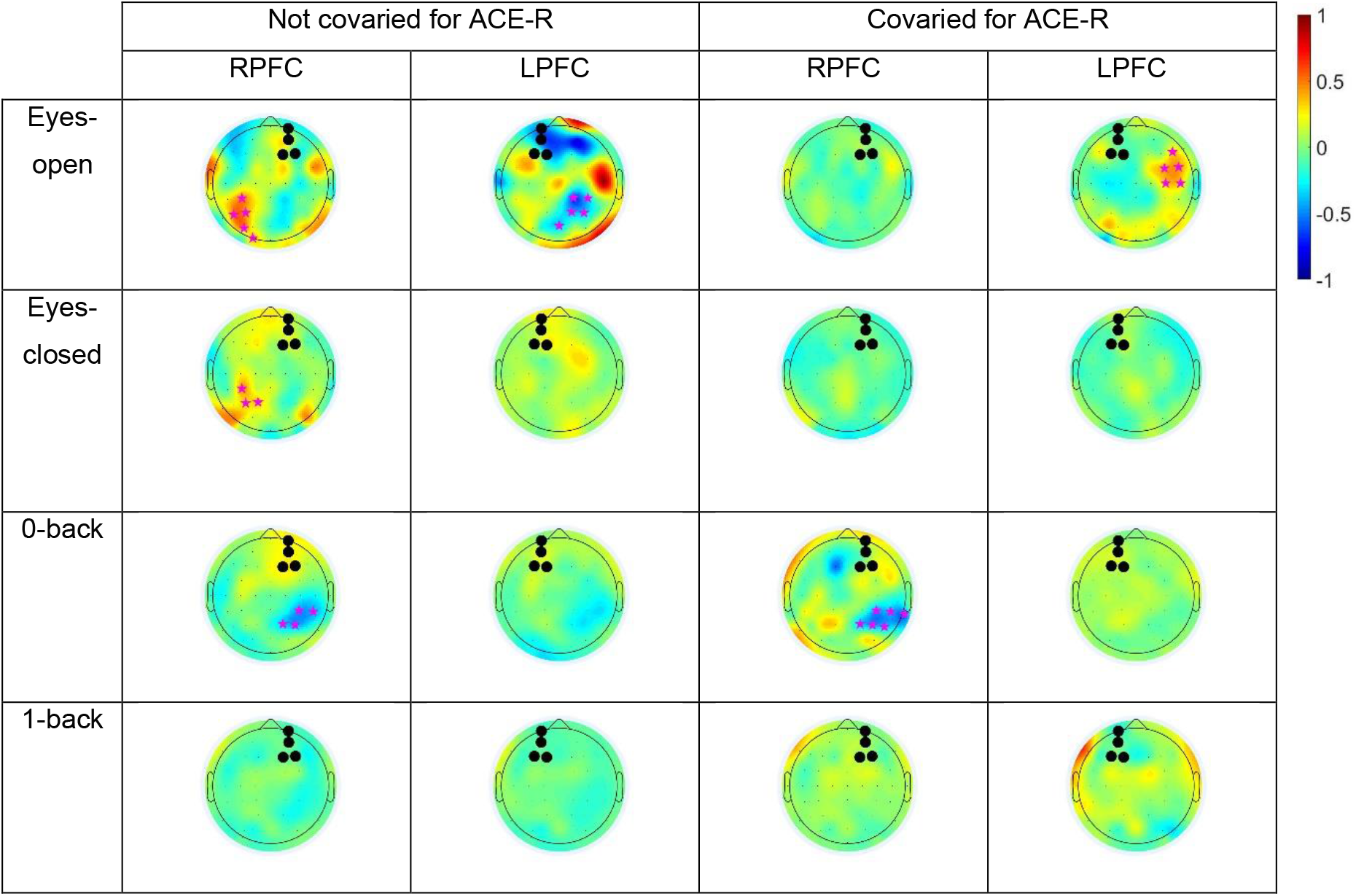
Topographic plots of regression coefficients from the parallel and orthogonalized partial least squares multiblock regression model in the theta band. The seed electrodes are shown with filled black circles. Colour bar represents regression coefficients.

**Table 2.**
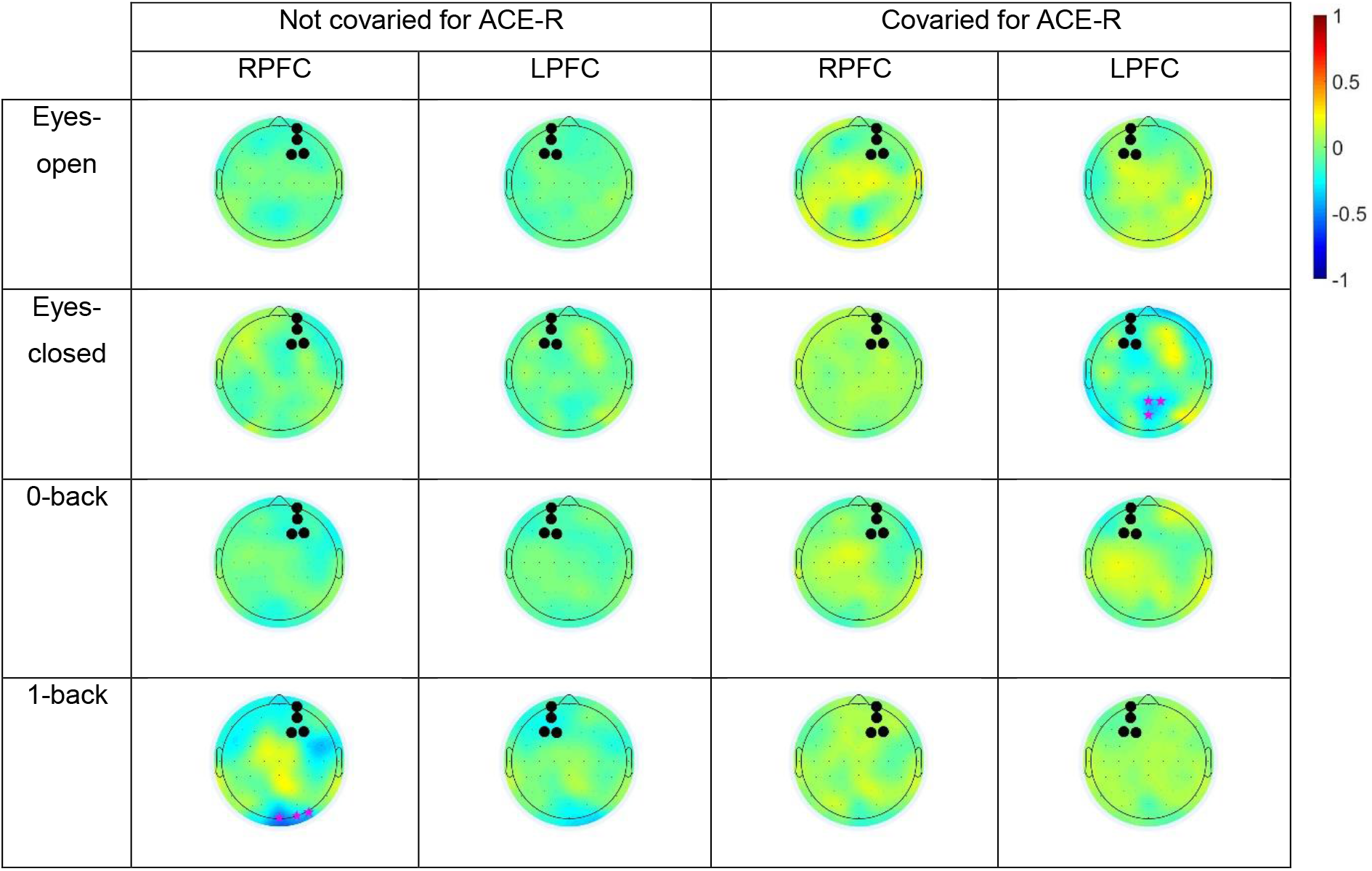
Topographic plots of regression coefficients from the parallel and orthogonalized partial least squares multiblock regression model in the alpha band. The seed electrodes are shown with filled black circles. Colour bar represent regression coefficients.

